# Sensing of extracellular ATP via P2RX7 drives lung tumor growth through regulatory T cell suppressive function

**DOI:** 10.1101/2025.05.03.652022

**Authors:** Igor Santiago-Carvalho, Ronaldo Francisco, Bruna de Gois Macedo, Caio Loureiro Salgado, Carly R. Stoll, Marcos Pinheiro Cione, Emily White, Tyler Johnston, Chloe Liliana Leff, Ildefonso Silva, Fabio Carvalho de Souza, Maria Regina D’Império Lima, Jessica Naomi Lancaster, Henrique Borges da Silva

## Abstract

Lung cancer is the leading cause of cancer-related deaths worldwide and, despite treatment advances, immune suppression remains an obstacle to effective therapy. Effector CD4^+^ T cells (CD4^+^ Teffs) are critical for antitumor immunity, but their function is often inhibited by regulatory T cells (Tregs), which accumulate in lung tumors and perform suppressive functions through multiple mechanisms. This suppression leads to tumor progression and poor patient outcomes. However, the mechanisms underlying Treg-mediated suppression are not fully understood. Here, we identify the extracellular ATP receptor P2RX7 as a key regulator of Treg function in lung tumors. Using a murine lung cancer model induced by Lewis lung carcinoma cells, we demonstrate that P2RX7 enhances the suppressive capacity of tumor-infiltrating Tregs, promoting tumor growth. In T cell-specific P2RX7-KO mice, reduced Treg infiltration was accompanied by increased CD4^+^ Teff accumulation and improved tumor control. Treg-specific P2RX7-KO mice exhibit reduced tumor growth, confirming a cell-intrinsic role of P2RX7 in Tregs. Suppression assays revealed that tumor-infiltrating WT Tregs have greater suppressive activity compared to P2RX7-KO Tregs, which failed to inhibit type 1 and Tfh-like responses. This was associated with increased tumor-specific IgG production by lung B cells in P2RX7-KO mice. We also observed that WT Tregs express higher levels of the immunosuppressive surface molecule CTLA-4 when compared to P2RX7-KO Tregs. In summary, we show that P2RX7 expression on Tregs is essential for their suppressive function in lung cancer, and targeting of P2RX7 may constitute a novel strategy to improve lung cancer treatment by alleviating Treg-mediated immune suppression.

## INTRODUCTION

Lung cancer remains the leading cause of cancer-related deaths worldwide, accounting for over 1.8 million deaths annually worldwide (1). Despite recent advances in early diagnosis and the development of immunotherapies, the prognosis for patients with lung cancer, especially non-small cell lung cancer (NSCLC), remains poor (2). A major challenge to successful therapies is the immunosuppressive environment established within the tumor, which hampers effective antitumor immune responses (3).

CD4^+^ T cells are key players in the immune response against lung tumors (4). These cells orchestrate antitumor immunity through many mechanisms, including direct cytotoxicity against tumor cells, support for cytotoxic CD8^+^ T cells, and collaboration with B cells to induce the production of tumor-specific antibodies (5, 6). In both human and murine lung tumors, effector CD4^+^ T cells (CD4^+^ Teffs) have been shown to contribute to tumor control and to enhance the quality and magnitude of cytotoxic responses (7). Furthermore, CD4^+^ T cell help is critical for the formation and maintenance of intratumoral tertiary lymphoid structures (TLS), which are associated with improved prognosis and better responses to immunotherapy (8). However, despite their importance, the antitumor function of CD4^+^ Teffs is often undermined within the tumor microenvironment (TME) (4).

A major contributor to Teff dysfunction is the accumulation and suppressive activity of FOXP3^+^ regulatory CD4^+^ T cells (Tregs) (9). Tregs are enriched in lung tumors and inhibit Teff proliferation, cytokine production, and cytotoxic activity (10, 11). Their presence is frequently associated with worse clinical outcomes and resistance to immune checkpoint blockade (12). Tregs can limit CD4^+^ Teff-mediated help for B cells and CD8^+^ T cells, dampening the immune system’s ability to eliminate tumor cells (13,14). Given their central role in immunosuppression, understanding the mechanisms that govern Treg accumulation and function in the lung TME is essential for developing strategies to restore effective antitumor immunity.

Several factors have been implicated in driving Treg function in tumors, such as cytokines, metabolic cues, and engagement of co-inhibitory receptors (15). However, the role of damage-associated molecular patterns (DAMPs), such as extracellular ATP (eATP), in shaping Treg responses in cancer remains incompletely understood (16). eATP is often released in inflamed or damaged tissues, including tumors, and is sensed by immune cells through purinergic receptors (17, 18). Among these receptors, P2RX7 has emerged as a key regulator of immune cell fate and function in inflammatory and infectious settings (19). In Teffs, P2RX7 signaling has been linked to cell survival, tissue residency, and effector differentiation (20, 21, 22). However, whether and how P2RX7 contributes to the immunosuppressive activity of Tregs in the context of lung cancer has not been investigated.

In this study, we identified P2RX7 as a critical regulator of Treg function in lung tumors. Using human NSCLC public data and murine models of lung cancer, we found that P2RX7 promotes the suppressive capacity of tumor-infiltrating Tregs and favors their accumulation in the TME. CD4^+^ T cell-specific P2RX7-KO mice exhibit reduced tumor growth, accompanied by a significant increase in Teff infiltration and antitumor activity. Importantly, Treg-specific deletion of P2RX7 revealed an Treg-intrinsic role of this receptor in enhancing Treg-mediated immunosuppression. We also found that lung P2RX7-KO Tregs fail to upregulate the key suppressive co-inhibitory receptor CTLA-4, and show diminished ability to suppress both Th1 and Tfh-like responses. Together, our findings reveal a novel role for P2RX7 in promoting Treg-mediated suppression in lung cancer and suggest that targeting this pathway may shift the balance between Tregs and Teffs, unlocking antitumor immunity.

## MATERIALS AND METHODS

### Single cell RNA-seq

Single-cell RNA sequencing (scRNA-seq) data from 42 samples of patients with NSCLC were obtained from the publicly available GEO NCBI repository (GSE148071) (23). Downstream analyses were performed using the Seurat v5 workflow in R. Each sample was individually filtered to exclude cells expressing fewer than 200 genes and 350 RNA features or exhibiting mitochondrial gene expression above the 95th percentile. Additionally, cells with gene counts above the 95th percentile were also removed. Data integration was performed using Harmony, implemented within Seurat. After loading the count matrices into Seurat objects, we applied NormalizeData, FindVariableFeatures, ScaleData, and RunPCA (n_dims = 50), followed by batch correction with RunHarmony. Clustering was performed by running FindNeighbors and FindClusters, and cell types were assigned using the human Lung v2 reference from Azimuth (https://azimuth.hubmapconsortium.org). Cluster annotations were further refined based on canonical marker gene expression. Differential gene expression analysis was conducted using Seurat’s FindAllMarkers function, with significance thresholds set at p-adjusted < 0.1, |log2 fold change| ≥ 1, and expression in at least 30% of cells.

### Spatial transcriptomics

10x Genomics Visium spatial transcriptomic data from 8 patients with NSCLC were obtained from the Human Tumor Atlas Network (HTAN) consortium (https://humantumoratlas.org). A total of 20 spatial transcriptomic slides were analyzed, with 2 to 4 tissue sections profiled per patient. For each patient, samples were collected from distinct tumor regions within the same lung lobe or from different lobes to capture intratumoral spatial heterogeneity. Raw data processing and downstream analyses were conducted in R using the Seurat package. Spot transcriptomes were filtered to exclude spots with fewer than 300 detected genes. Sampling areas with fewer than 20 high-quality spot transcriptomes were excluded from further analysis. Count matrices were normalized and scaled using NormalizeData and ScaleData functions with default parameters. Deconvolution analysis was performed using RCTD (24), as implemented in the spacexr package. Single-cell RNA-seq data served as the reference. For each sampling area, an RCTD object was created by combining the single-cell reference and the corresponding SpatialRNA object. RCTD analysis was run using the run.RCTD function with the doublet_mode parameter set to "full".

### Survival probability analysis

Survival data from over 1,000 NSCLC patients were analyzed using the Kaplan-Meier Plotter tool (https://kmplot.com/analysis). Overall survival was evaluated based on *P2RX7* mRNA expression levels. Patients were stratified into “high” and “low” expression groups according to their *P2RX7* median value. Kaplan-Meier survival curves were generated, and statistical significance was determined using the long-rank test.

### Mice

Male and female mice, aged 6-8 weeks, with a specific-pathogen-free status, were used in this study. Mice were housed in the animal facilities of The Department of Immunology at Mayo Clinic Arizona. All mice were randomly assigned to experimental groups. The following mouse strains were utilized: C57BL/6, P2RX7-KO, RAG2-KO, CD4-cre, CD4-cre *P2rx7^flox/flex^*, Foxp3 eGFP-Cre-ERT2 and Foxp3 eGFP-Cre-ERT2 *P2rx7^flox/flox^*. All strains were obtained from The Jackson Laboratory, except for the *P2rx7*^flox/flox^ strain which was obtained from Matyas Sandor (University of Wisconsin). In all experiments, mice were sacrificed by isoflurane overdose inhalation. All experimental procedures were conducted in accordance with institutional guidelines and were approved by the institutional Animal Care and Use Committee (IACUC - A00005542-20-R23).

### Cell lines and tumor injection

The Lewis lung carcinoma (LLC) cell line (ATCC), both GFP-expressing (LLC-GFP) and non-GFP (LLC) variants, was used for lung cancer murine experiments. LLC cells were cultured in complete DMEM (Dulbecco’s Modified Eagle Medium - Gibco) supplemented with 10% fetal bovine serum (FBS – Cardinal Healthcare), 1% penicillin-streptomycin, and 1% L-glutamine at 37°C in a 5% CO₂ atmosphere. Cells were maintained in an exponential growth phase and were used for injections after confirming proper growth and morphology. LLC-GFP cells (5x10^5^) were intravenously injected into experimental mice to establish lung metastasis (25), while LLC cells were used in some experiments to assess tumor growth without GFP labeling.

### CD4^+^ T cell purification and adoptive transfer experiments

For single adoptive transfer experiments, splenocytes were isolated from C57BL/6 (CD45.1, WT) and T-cell-specific P2RX7-KO (CD45.2) mice. Naïve CD4^+^ T cells were purified from splenocytes by negative selection using the EasySep™ Mouse Naïve CD4^+^ T Cell Isolation Kit (STEMCELL Technologies), following the manufacturer’s instructions. The negative fraction, containing splenocytes depleted of CD4^+^ T cells, was washed with 1x PBS and transferred into RAG2-KO mice. Mice were randomly assigned into two groups, receiving either WT or P2RX7-KO naive CD4^+^ T cells. After 24 hours, both groups were injected intravenously with LLC-GFP tumor cells. Lungs were harvested at day 30 post-LLC injection (p.i.) for histological and flow cytometry analyses.

For co-adoptive transfer experiments, splenocytes from WT (CD45.1) and P2RX7-KO (CD45.2) mice were isolated and co-transferred into RAG2-KO mice. At 24 hours post-transfer, recipients were injected with 5x10^5^ LLC-GFP cells. Lungs were collected on day 30 p.i. for flow cytometry analysis.

### Administration of tamoxifen in mice

On day 5, 7, 9 and 11 p.i., Foxp3 eGFP-Cre-ERT2 (WT) and Foxp3 eGFP-Cre-ERT2 *P2rx7^flox/flox^* mice were treated intraperitonially with 1 mg of tamoxifen (Sigma-Aldrich) diluted in sunflower oil (vehicle) (26).

### Tissue processing for flow cytometry and ELISA

On day 30 p.i., lungs were harvested, and the lobes were processed and digested with collagenase type IV (0.5 mg/mL) (Gibco) at 37°C for 40 minutes under agitation (200 rpm) (22). Mediastinal lymph nodes (medLNs) and spleens were mechanically processed using cell strainers (Corning). Lung homogenates were separated for IgG measurement by ELISA. The cell suspension obtained from the tissues was homogenized, filtered through cell filters, and incubated with ACK lysis buffer (made in house) at room temperature for 2 minutes to deplete erythrocytes. The cell suspensions were washed with PBS containing 10% FBS. Afterward, the cell suspension was centrifuged at 1,200 rpm for 5 minutes and resuspended in FACS buffer until the time of staining.

### Flow cytometry analysis

Cell suspensions from the lung, medLN and spleen were stained with fluorochrome-conjugated monoclonal antibodies (see **Supplementary Table 1**) diluted in FACS buffer and incubated at room temperature (RT) for 40 minutes. After staining, cells were washed with FACS buffer and then prepared for intracellular IFN-γ staining (22). For this, lung cells were incubated with eBioscience™ Cell Stimulation Cocktail plus protein transport inhibitors (Invitrogen - 2 μM) for 4 hours at 37°C in 5% CO2. After incubation, cells were fixed and permeabilized using the BD Cytofix/Cytoperm™ Fixation/Permeabilization Kit (BD Biosciences) and then washed with the buffer provided in the kit. ViaDye™ Red Fixable Viability Dye Kit (Cytek Biosciences) was used to identify dead cells. Samples were analyzed using multiparametric flow cytometry with a 27-parameter panel on Cytek Aurora (Cytek Biosciences) flow cytometer. Data was processed for conventional analysis and cluster distribution through t-distributed stochastic neighbor embedding (tSNE) using FlowJo software (BD Biosciences).

### Histological analysis

The upper left lung lobes were collected, rinsed in PBS, and fixed in 4% paraformaldehyde (PFA) for 24 hours at 4°C. Fixed tissues were then embedded in paraffin, sectioned into 4-5 μm slices, and stained with hematoxylin and eosin (H&E) to evaluate tissue pathology. Stained sections were visualized under a light microscope and representative imagens were acquired for analysis.

### Fluorescence microscopy

PFA-fixed paraffin-embedded samples were deparaffinized and subjected to antigen retrieval by steaming for 20 minutes in 1X Citrate Buffer (Diagnostic Biosystems). Next, sections were permeabilized with 2% Triton-X 100 in 1% bovine serum albumin (BSA) in PBS for 30 minutes, followed by blocking in 5% BSA in PBS for 30 minutes at RT. Sections were stained for 48 hours at 4°C in a humidity chamber in the dark with the following primary antibodies diluted in BSA blocking buffer: rabbit polyclonal anti-CD4 (1:50, Abcam) and mouse anti-Ep-CAM (1:100, Invitrogen) antibodies. Sections were washed three times with 0.1% Tween-20 in PBS before incubation for 2 hours at RT in the dark with goat anti-rabbit IgG AF546 (Invitrogen A11035) and goat anti-mouse IgG AF488 (Invitrogen A11001) secondary antibodies diluted (1:200) in BSA blocking buffer. Slides were counterstained with DAPI for 15 minutes, washed thrice, and then mounted with ProLong™ Gold Antifade Mountant (Thermo Fisher Scientific) and secured with a coverslip. Sections were imaged using LSM 800 confocal microscope (Zeiss) equipped with Plan-Neofluar 20× objective lens (Zeiss, NA 0.4).

### Suppression assays

Tregs (Foxp3/GFP^+^CD25^+^CD4^+^ T cells) were sorted from the lung of WT and Treg-specific P2RX7-KO mice at day 30 p.i. using MA900 Multi-Application Cell Sorter (Sony Biotechnology). Sorted lung Tregs were plated with pre-activated WT CD4^+^ and CD8^+^ T cells isolated from the spleen with appropriated columns at 0:1, 1:5 and 1:20 and Treg/Teff ratios (STEMCELL Technologies) and stimulated *in vitro* with anti-CD3/CD28 antibodies. Teffs were labeled with CellTrace™ Violet (Thermo Fisher Scientific), and Tregs were labeled with CFSE (Carboxyfluorescein succinimidyl ester), to allow tracking of proliferation and identification of distinct populations within the same well. After 4 days in culture, cells were collected, washed with FACS buffer, and stained for flow cytometry analysis. The percentage of suppression was calculated using the following formula:

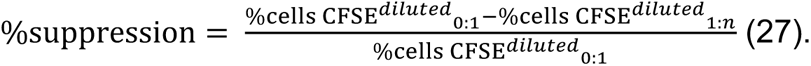

### *In vitro* Treg activation and treatments

Splenic Tregs were isolated using EasySep™ Mouse CD4^+^CD25^+^ Regulatory T Cell Isolation Kit II (STEMCELL Technologies), following the manufacturer’s protocol. Cells were resuspended in RPMI medium supplemented with 10% of FBS, 1% L-glutamine, 1% sodium pyruvate, and 1% penicillin-streptomycin, then plated (5x10^4^) in 96-well plates pre-coated with anti-CD3 and anti-CD28 antibodies for *in vitro* activation. Interleukin-2 (mouse IL-2, 10 ng/ml) was added daily to the culture. Some wells received the P2RX7 agonist HEI3090 (Axon Medchem, 50μM) at the time of plating (28). After 72 hours, cells were washed with FACS buffer and stained for flow cytometry analysis.

### Calcium influx assays

For assessment of calcium influx, cells were stained with surface markers, then incubated for 1 h at 37 °C with 1 μM Fluo-4-AM-FITC (ION Biosciences) and then washed with HEPES buffered saline (made in house) (29). Cells were analyzed by kinetic (Time vs Fluo-4-AM-FITC) flow cytometry, with a baseline reading being determined for 5 min followed by ionomycin (STEMCELL Technologies) addition, with the analysis continued for the remaining 20 min. Subsequently, the P2RX7 agonist HEI3090 (Axon Medchem – 50 μM) was added, and cells were acquired for an additional 10 minutes. After this period, the P2RX7 inhibitor A438079 (MedChemExpress – 100 μM) was added (30), and cells were analyzed for another 10 minutes. The entire acquisition was performed at low speed and continuously, with the sample tube being removed for cell treatments without stopping acquisition. Flow cytometric analysis was performed using a FACSymphony™ (BD Biosciences) and data were analyzed with FlowJo software (BD Biosciences).

### Anti-tumor IgG quantification

Tumor-specific IgG antibodies were quantified using an indirect ELISA approach. LLC tumor cells were lysed using RIPA buffer, and the resulting lysates were used to coat high-binding flat-bottom 96-well plates for 24 hours at 4°C. The following day, standard indirect ELISA procedures were performed using serum and lung homogenate collected from WT and P2RX7-KO mice on day 30 p.i. (31). All antibodies and reagents used in this protocol are listed in the Key Resources Table. Optical density was measured at 450 nm using a Synergy HTX Multi-Mode plate reader (BioTek).

### Statistical analysis

Specific statistical tests applied to each experiment are detailed in the respective figure legends. All analyses were conducted using GraphPad Prism 10.4.0 software. Data are presented as Mean values with standard deviation (SD) shown as error bars. For comparisons between two groups, Paired or Unpaired T-tests were used. Differences between groups were considered significant when p < 0.05 (*), p < 0.01 (**) or p < 0.001 (***).

### Data availability statement

The data generated in this study are available upon request from the corresponding authors.

## RESULTS

### P2RX7 is highly expressed in tumor-infiltrating T cells of lung cancer patients

To investigate the association between *P2RX7* expression and lung cancer outcomes, we analyzed clinical datasets from The Cancer Genome Atlas (TCGA). Patients with higher *P2RX7* mRNA expression exhibited significantly poorer overall survival compared to those with lower expression levels (**Fig. 1A**). Next, the *P2RX7* expression at the cellular level was evaluated using the scRNA-seq data from the TME of NSCLC patients. After quality control, 90,406 high-quality cells were retained from 42 advanced-stage NSCLC samples. These cells are clustered into nine major groups, comprising structural cells (epithelial, endothelial, and fibroblasts) and immune cells (macrophages, dendritic cells, B cells, and T cells) (**Fig. 1B**). *P2RX7* expression was detected across all clusters, with the highest levels found in T cells (**Fig. 1C**). Further analysis of CD4^+^ T cell clusters revealed three main populations: proliferating, central memory (TCM), and Tregs (**Fig. 1D**). Among these, Tregs expressed the highest level of *P2RX7* (**Fig. 1E**).

**Fig. 1.**
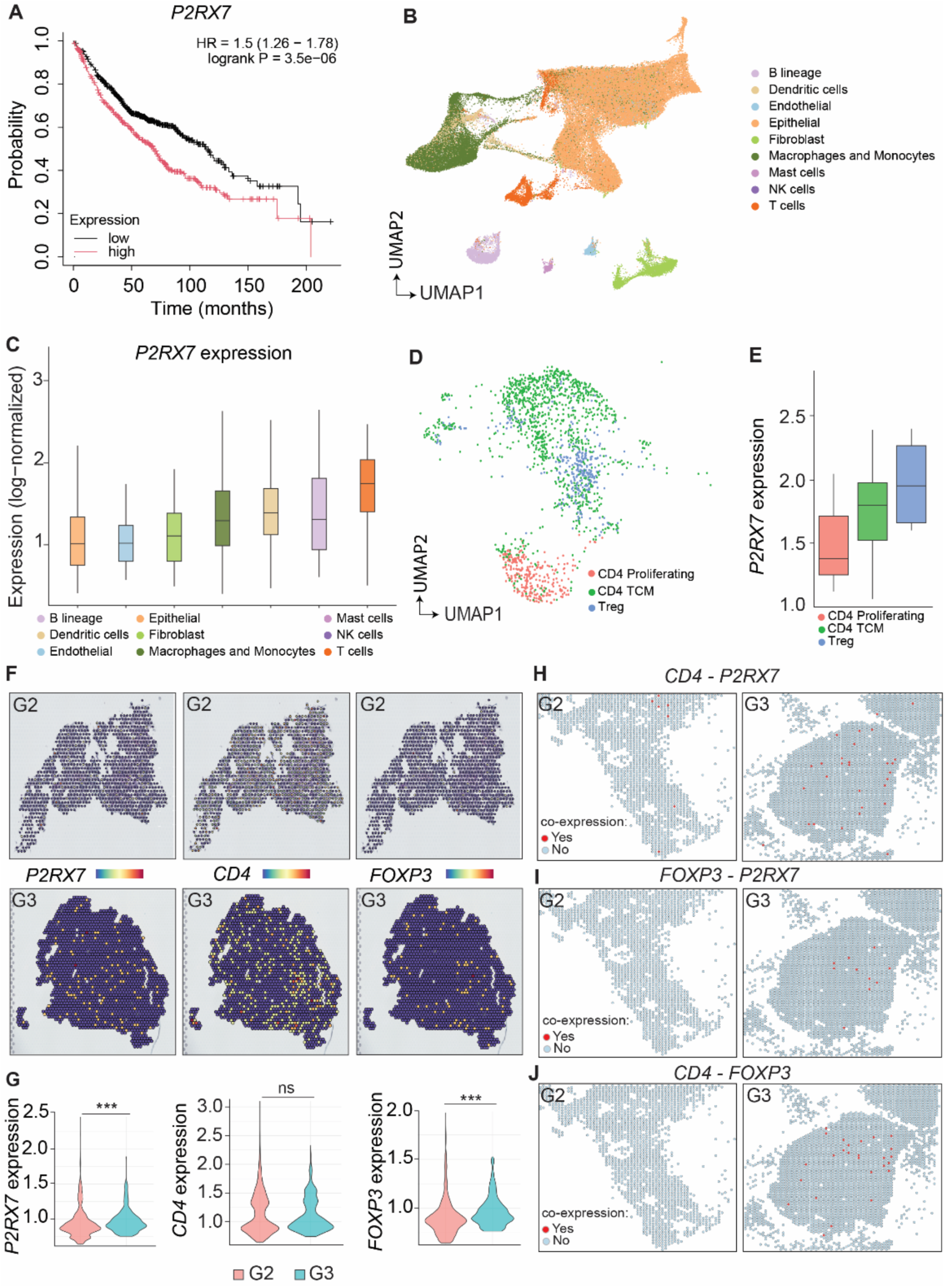
*P2RX7* expression on immune cells in lung cancer patients. (**A**) Kaplan-Meier curves showing the overall survival values related to P2RX7 expression in TCGA database. (**B**) UMAP projection of 9 clusters of immune and non-immune cells in human lungs with cancer. (**C**) Boxplot showing the average expression of *P2RX7* in immune and non-immune cell clusters in human lungs with cancer. (**D**) UMAP showing the subclusters of CD4^+^ T cells in human lungs with cancer. (**E**) Boxplot showing the average expression of *P2RX7* in CD4^+^ T cells subclusters in human lungs with cancer. (**F**) Spatial images showing the spots expressing *P2RX*7, *CD4* and *FOXP3* in lung cancer grade 2 (G2) and grade 3 (G3) patients. (**G**) Violin plots showing the average expression of *P2RX7*, *CD4* and *FOXP3* in lung cancer G2 and G3 patients. (**H**) Spatial images showing the location of spots co-expressing *CD4* and *P2RX7* G2 and G3 lung cancer patients. (**I**) Spatial images showing the location of spots co-expressing *FOXP3* and *P2RX7* in G2 and G3 lung cancer patients. (**J**) Spatial images showing the location of spots co-expressing *CD4* and *FOXP3* in G2 and G3 lung cancer patients. Spatial images are representative sections from two patients. Differences in *P2RX7* expression between tumor grades were assessed using the Wilcoxon rank-sum test. p-values were calculated and displayed on the plot.

We further analyzed spatial transcriptomics data from NSCLC patients to assess the expression patterns of *P2RX7*, *CD4*, and *FOXP3* across distinct regions of tumor tissue (24). A spatial trend in *P2RX7* expression was observed in relation to tumor grade. Multiple *P2RX7*-positive spots were detected in both grade 2 (G2) and grade 3 (G3) tumors, with higher expression levels observed in G3 samples (Fig. 1F–G). *CD4* expression was detected in both tumor grades; however, no significant differences in overall CD4 expression were observed between G2 and G3 tumors. Similar to *P2RX7*, *FOXP3* expression appeared higher in G3 compared to G2 tumors (Fig. 1F–G). Both *P2RX7* and *FOXP3* showed increased expression levels in samples from patients with higher tumor grade. Finally, we analyzed the co-expression of these genes and observed a greater number of regions co-expressing *CD4 - P2RX7*, *FOXP3 - P2RX7* and *CD4 - FOXP3* in G3 patients (**Fig. 1H-J**), when compared to G2 patients. Together, these data support the notion that CD4^+^ T cell-intrinsic P2RX7 expression in Tregs may serve as a negative prognostic biomarker in lung cancer.

### Tregs exhibit higher P2RX7 expression than CD4^+^ Teffs in response to murine lung cancer

To investigate the *in vivo* role of P2RX7 in T cell responses to lung cancer, we used an experimental model induced by the LLC cell line (**Fig. 2A**). To facilitate tumor visualization in the lung, we used GFP-expressing LLC cells (LLC-GFP). Cancer cells were injected intravenously (i.v.), and 30 days p.i., we harvested the lungs and medLNs for analysis. GFP^+^ tumor nodules were macroscopically visible through the lung tissue (**Fig. 2B, left**). Histological analysis showed widespread tumor development beyond the visible nodules, with areas of immune cell infiltration (**Fig. 2B, right**), indicating a robust ongoing anti-tumor immune response by day 30 p.i. We next analyzed activated CD4^+^ T cells (CD44^+^) in the lungs and medLNs of tumor-bearing mice (gating strategy in **Fig. S1**). In both organs, P2RX7 expression was heterogeneous, with some cells exhibiting high levels and others exhibiting low levels of the receptor (**Fig. 2C)**. Next, we separated CD44^+^CD4^+^ T cells into CD4^+^ Teffs and Tregs and compared P2RX7 expression between these populations in both organs. Our data shows that among CD44^+^CD4^+^ T cells, Tregs express higher levels of P2RX7 than CD4^+^ Teffs (**Fig. 2D-E**). These findings indicate that lung tumors induce the accumulation of P2RX7-expressing CD44^+^ CD4^+^ T cells, with Tregs expressing higher levels of this receptor.

**Fig. 2.**
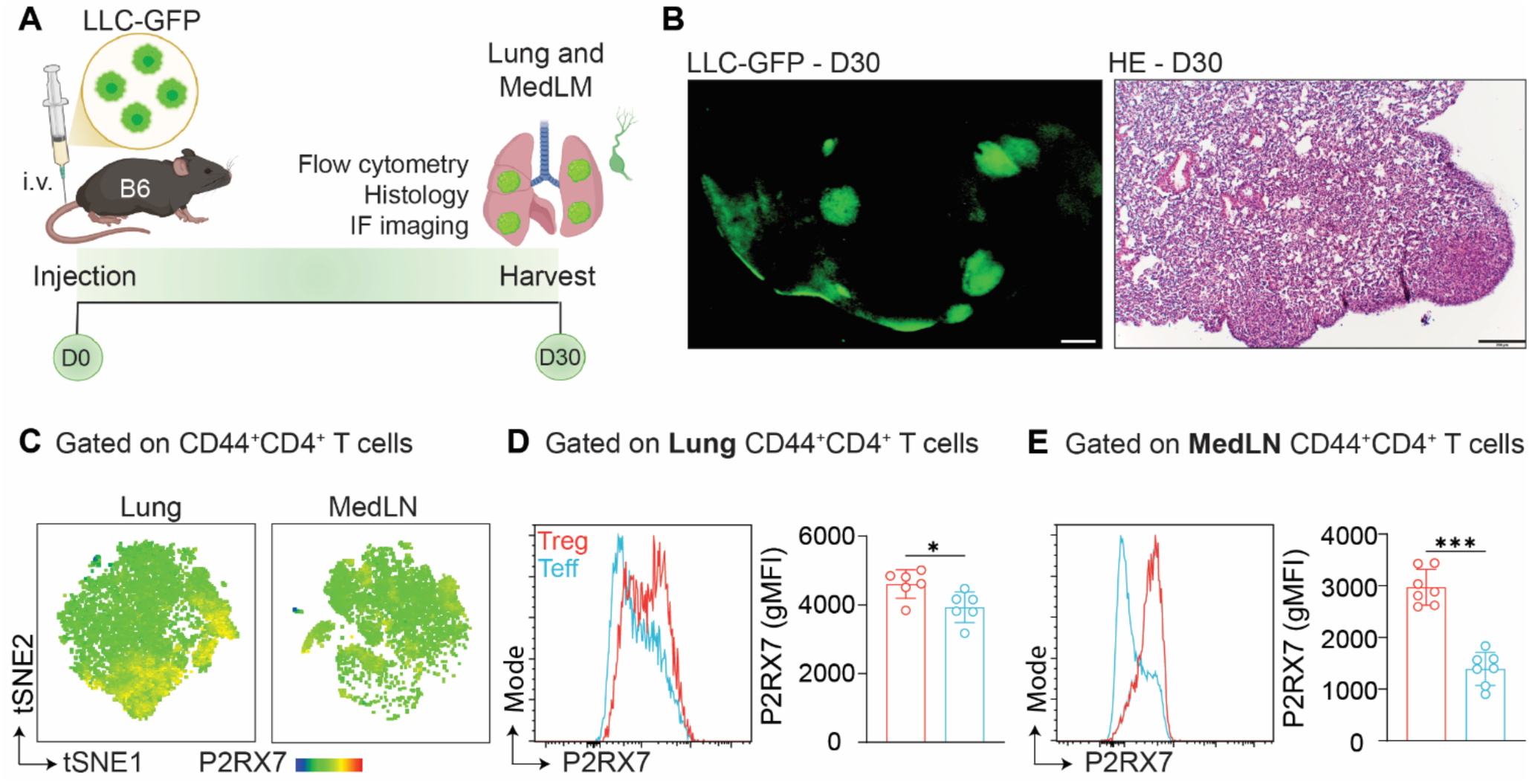
P2RX7 expression on CD4^+^ T cells in lung cancer metastasis experimental model. C57BL/6 mice were injected i.v. with LLC-GFP cells and lungs and medLNs were collected 30 days p.i. (**A**) Schematic illustration showing the lung cancer experimental protocol (BioRender.com). (**B**) Left: Representative fluorescence macroscopy of the left upper lung lobes with tumors (20 μm). Right: Representative section of the left upper lung lobes stained with H&E (200 μm). (**C**) tSNE showing the P2RX7 expression on CD44^+^CD4^+^ T cells in lung cancer. (**D-E**) Left: histograms of P2RX7 expression on CD44^+^CD4^+^ Tregs and Teffs. Right: average geometric mean fluorescence intensity (gMFI) of P2RX7 on CD44^+^CD4^+^ Tregs and Teffs. Data from 2-3 independent experiments. Data shown as means ± SD. *p < 0.05, **p < 0.01, ***p < 0.001. Statistical significance was determined by unpaired t tests.

### T-cell-specific P2RX7 expression permits tumor growth in the lungs

To investigate the role of P2RX7 specifically on T cells in lung cancer, we injected LLC-GFP cells into WT (CD4-cre) and T cell-specific P2RX7-KO mice (CD4-cre *P2rx7^fl/fl^*) mice (**Fig. 3A**). Over a 30-day period, we monitored body weight and observed that, while WT mice experienced weight loss or lack of weight gain, T cell-specific P2RX7-KO mice continued to gain weight over time (**Fig. 3B**). In line with apparent increased disease development, WT mice exhibited heavier lungs compared to P2RX7-KO mice (**Fig. 3C**). Macroscopic fluorescence imaging revealed larger and more disseminated GFP^+^ tumor nodules in WT than in P2RX7-KO lungs (**Fig. 3D and S2A-B**). Flow cytometry confirmed a higher number of LLC-GFP cells in WT lungs (**Fig. 3E and Fig. S2C**). Histological analysis showed extensive tumor areas in WT lungs, whereas P2RX7-KO lungs displayed smaller tumor regions, contrasting with a pronounced immune cell infiltrate in P2RX7-KO LLC-inoculated lungs (**Fig. 3F**).

**Fig. 3.**
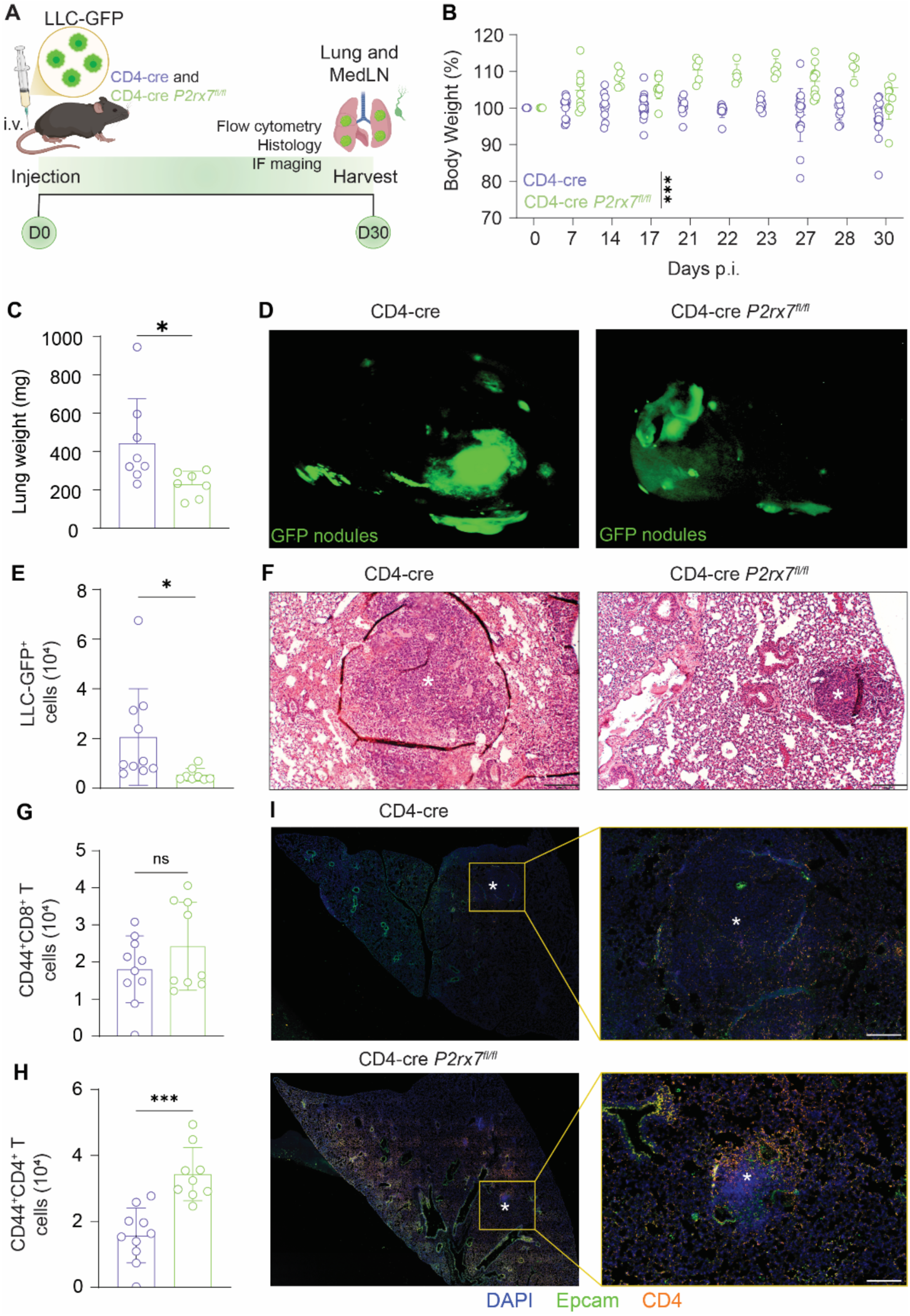
Effects of T-cell intrinsic P2RX7 in lung cancer experimental model. WT (CD4-cre) and T cell-P2RX7-KO (CD4-cre *P2rx7^fl/fl^*) mice were injected i.v. with LLC-GFP cells and lungs were collected on day 30 p.i. (**A**) Schematic illustration showing the lung cancer experimental protocol (BioRender.com). (**B**) Average body weights (percentages related to day 0). (**C**) Lung weight values on day 30 p.i. (**D**) Representative fluorescence macroscopy of the left upper lung lobes with tumors (20 μm). (**E**) Average numbers of GFP^+^ LLC cells in the lungs. (**F**) Representative section of the left upper lung lobes stained with H&E (200 μm). (**G**) Average numbers of CD44^+^CD8^+^ T cells in the lungs. (**H**) Average numbers of CD44^+^CD4^+^ T cells in the lungs. (**I**) Representative confocal images of DAPI (blue), Epcam (green) and CD4 (orange) staining in the lung tissues (50 μm). Data from 2-3 independent experiments. Data shown as means ± SD. *p < 0.05, **p < 0.01, ***p < 0.001. Statistical significance was determined by unpaired t tests.

Next, we analyzed the phenotypic characteristics of lung-infiltrating T cells. The number of CD44^+^CD8^+^ T cells was comparable between WT and P2RX7-KO mice (**Fig. 3G**). However, P2RX7-KO lungs contained higher numbers of CD44^+^CD4^+^ T cells (**Fig. 3H**). This was further confirmed using confocal microscopy, showing that while in WT mice CD4^+^ T cells were less abundant and sparsely distributed, P2RX7-KO mice exhibited an increased CD4^+^ T cell numbers across the lung tissue (**Fig. 3I**). Collectively, our findings suggest that T-cell intrinsic P2RX7 allows lung tumor growth while limiting the accumulation of CD4^+^ Teffs within the lung tissue.

### T-cell Intrinsic P2RX7 favors Treg accumulation in lung tumors

To assess the T cell-intrinsic impact of P2RX7 expression on T cells within a competitive setting, we transferred splenocytes from WT and T-cell-specific P2RX7-KO mice into RAG2-KO recipients, followed by LLC-GFP tumor injection (**Fig. 4A**). At day 30 p.i., we analyzed host T cell numbers and found a higher frequency of activated P2RX7-KO CD4^+^ T cells (CD45.2^+^) compared to WT (CD45.1^+^) in the lung, as well as in the medLNs and spleens (**Fig. 4B**). Unlike previous observations in T-cell-specific P2RX7-KO mice, CD8^+^ P2RX7-KO T cells also accumulated in greater numbers than WT counterparts in the lung (**Fig. S3A**).

**Fig. 4.**
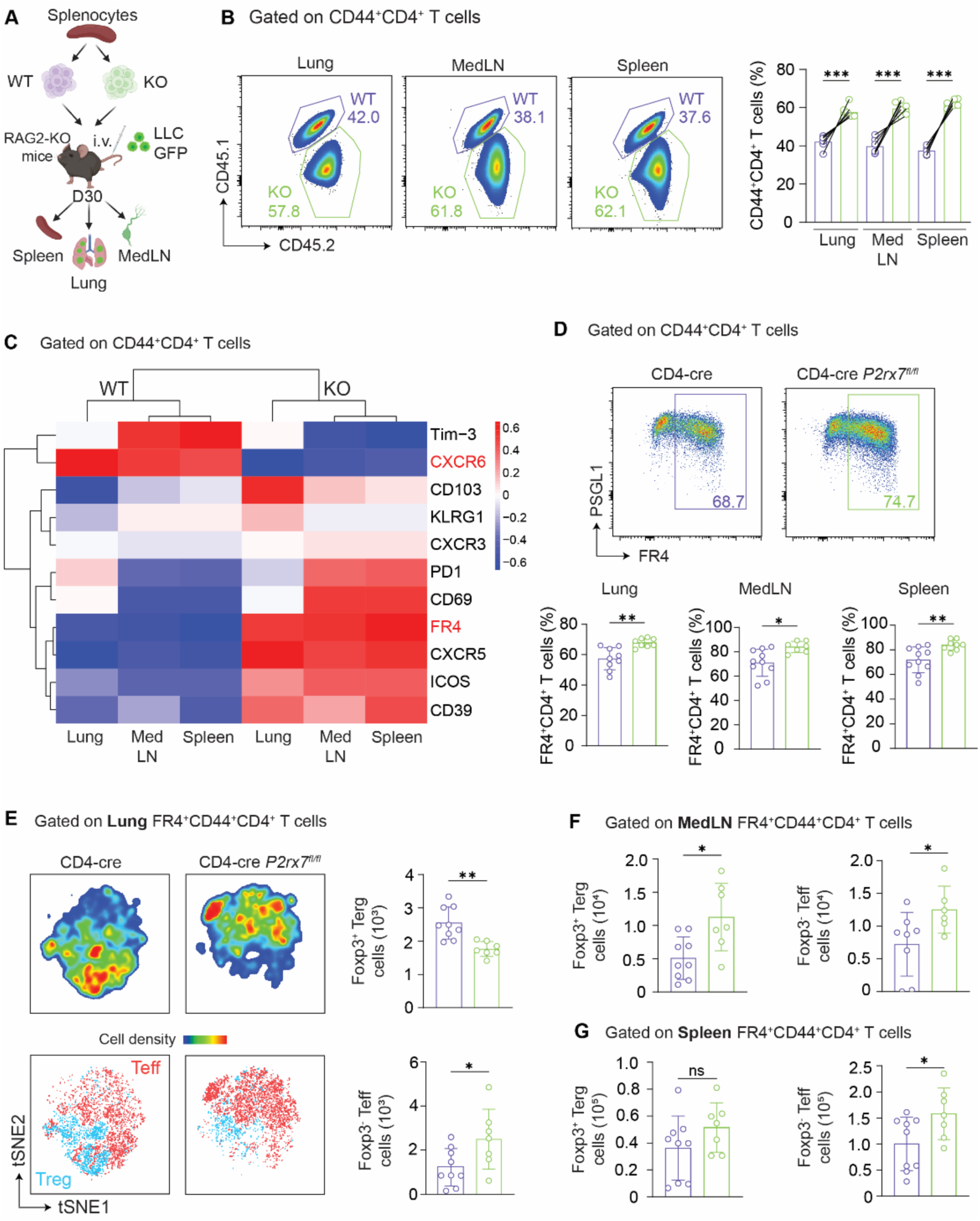
Effects of P2RX7 expression on CD4^+^ T cell accumulation in lung cancer experimental model. (**A-C)** RAG-KO mice were reconstituted with splenocytes from WT and T-cell-specific P2RX7-KO mice and injected i.v. with LLC-GFP cells. On day 30 p.i. the lungs, medLNs and spleens were collected. (**A**) Schematic illustration showing the lung cancer experimental protocol (BioRender.com). (**B**) Left: flow-cytometry plots of CD45.1 and CD45.2 expression on CD44^+^CD4^+^ T cells. Right: average frequencies and numbers of CD45.1 and CD42.2 CD44^+^CD4^+^ T cells. (**C**) Heatmap of gMFI between WT and P2RX7-KO CD44^+^CD4^+^ T cells in the lungs, medLNs and spleens. (**D-G**) WT (CD4-cre) and T cell-P2RX7-KO (CD4-cre *P2rx7^fl/fl^*) mice were injected i.v. with LLC-GFP cells and lungs were collected on day 30 p.i. (**D**) Top: flow-cytometry plots of PSGL1 and FR4 expression in the lungs. Bottom: average frequencies of FR4^+^CD44^+^CD4^+^ T cells in the lungs, medLNs and spleens. (**E**) Left: tSNE showing cell density and Foxp3 expression on FR4^+^CD44^+^CD4^+^ T cells in lung cancer. Right: Average numbers of CD44^+^CD4^+^ Tregs (Foxp3^+^) and Teffs (Foxp3^-^) in the lungs. (**F**) Average numbers of CD44^+^CD4^+^ Tregs (Foxp3^+^) and Teffs (Foxp3^-^) in the medLN. (**G**) Average numbers of CD44^+^CD4^+^ Tregs (Foxp3^+^) and Teffs (Foxp3^-^) in the spleen. Data from 2-3 independent experiments. Data shown as means ± SD. *p < 0.05, **p < 0.01, ***p < 0.001. Statistical significance was determined with paired (**B**) and unpaired t tests (**D-G**).

Next, we evaluated the phenotype of infiltrating T cells. While CD44^+^CD8^+^ T cells showed no phenotypic differences (**Fig. S3B**), WT CD44^+^CD4^+^ T cells exhibited higher expression of CXCR6 than P2RX7-KO cells in the lungs (**Fig. 4C and S3C**). Conversely, P2RX7-KO CD4^+^ T cells expressed higher levels of PD-1, CD69, CXCR5, ICOS, and CD39, with FR4 (Folate Receptor 4) being significantly increased across all organs analyzed (**Fig. 4C and S3C**). The frequency of FR4^+^CD4^+^ T cells was also slightly, but significantly higher in T-cell-specific P2RX7-KO mice than in WT controls in the lungs, medLNs, and spleens (**Fig. 4D**).

Since FR4 is expressed in both CD4^+^ Teffs and Tregs (32, 33), we distinguished these subsets using Foxp3 expression. T cell-specific P2RX7-KO mice exhibited an increased number of CD4^+^ Teffs in the lungs compared to WT, while Tregs were more abundant in WT lungs (**Fig. 4E**). In the medLNs of P2RX7-KO mice, both Tregs and CD4^+^ Teff numbers were increased (**Fig. 4F**). In the spleens, only P2RX7-KO Teff numbers were higher than WT ones (**Fig. 4G**). Taken together, our data indicates that T-cell-intrinsic P2RX7 expression supports Treg accumulation while limiting CD4^+^ Teff accumulation in tumors.

### eATP-P2RX7 axis on CD4^+^ T cells promotes Treg accumulation and allows tumor growth in the lungs

To evaluate whether P2RX7 expression in CD4^+^ T cells affects tumor growth, we used a single transfer model into RAG2-KO mice (**Fig. S4A**). In this single transfer model, P2RX7-KO CD4^+^ T cells prevented tumor growth or disease, differently from WT CD4^+^ T cells which allowed substantial weight loss and tumor development (**Fig. S4B-D**). We also observed higher numbers of WT Tregs compared to P2RX7-KO Tregs in both the lungs and medLNs (**Fig. S4E)**. In contrast, P2RX7-KO CD4^+^ Teffs increased only in the medLNs (**Fig. S4F**).

### Treg-intrinsic P2RX7 promotes immunosuppression favoring tumor growth in the lung

To evaluate the intrinsic effects of P2RX7 in Tregs, we employed a Foxp3-cre tamoxifen-inducible P2RX7 ablation system (**Fig. 5A**) Tamoxifen-induced ablation of P2RX7 in Tregs led to weight gain in tumor-bearing mice, whereas WT mice exhibited weight stability or loss (**Fig. 5B**). Additionally, lighter lungs were observed in Treg-specific P2RX7-KO mice at day 30 p.i. (**Fig. 5C**), accompanied by a reduced tumor area compared to WT mice (**Fig. 5D**).

**Fig. 5.**
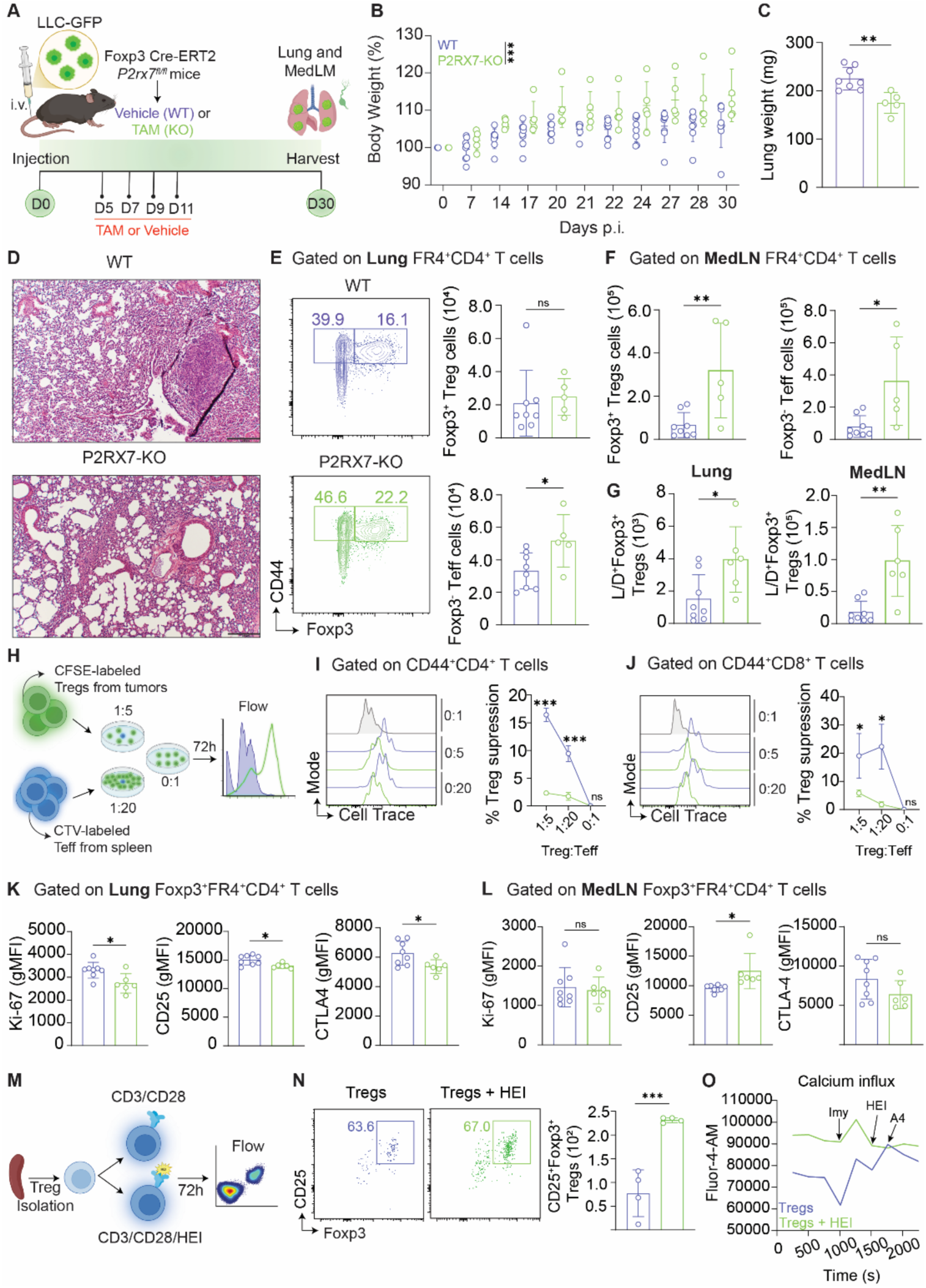
Effects of Treg-intrinsic P2RX7 on Treg suppressive activity and tumor growth in lung cancer experimental model. (**A-L**) WT (Foxp3 Cre-ERT2 *P2rx7^fl/fl^* - Vehicle) and Treg-specific P2RX7-KO (Foxp3 Cre-ERT2 *P2rx7 ^fl/fl^* – tamoxifen) mice were injected i.v. with LLC-GFP cells and lung were collected on day 30 p.i. (**A**) Schematic illustration showing the lung cancer experimental protocol (BioRender.com). (**B**) Average body weights (percentages related to day 0). (**C**) Lung weight values on day 30 p.i. (**D**) Representative section of the left upper lung lobes stained with H&E (200 μm). (**E**) Left: flow-cytometry plots of CD44 and Foxp3 expression in the lungs. Right: average numbers of Tregs (Foxp3^+^) and Teff (Foxp3^-^) CD44^+^CD4^+^ T cells in the lungs. (**F**) Average numbers of Tregs (Foxp3^+^) and Teff (Foxp3^-^) CD44^+^CD4^+^ T cells in the medLNs. (**G**) Average numbers of Live/dead^+^ Tregs (Foxp3^+^) in the lungs and medLNs. (**H**) Schematic illustration showing the suppression assay using WT and P2RX7-KO lung Tregs lung experimental protocol (BioRender.com). (**I**) Left: Histograms showing the Cell Trace staining on CD44^+^CD4^+^ T cells. Right: Average percentage of Treg suppression on CD44^+^CD4^+^ T cell proliferation. (**J**) Left: Histograms showing the Cell Trace staining on CD44^+^CD8^+^ T cells. Right: Average percentage of Treg suppression on CD44^+^CD8^+^ T cell proliferation. (**K**) Average geometric mean fluorescence intensity (gMFI) of Ki-67, CD25 and CTLA-4 on lung Tregs (Foxp3^+^). (**L**) Average geometric mean fluorescence intensity (gMFI) of Ki-67, CD25 and CTLA-4 on medLNs Tregs (Foxp3^+^). (**M**) Schematic illustration showing the *in vitro* activation of Tregs using P2RX7 agonist (HEI3090) or not (BioRender.com). (**N**) Left: flow-cytometry plots of CD25 and Foxp3 expression in in vitro activated Tregs. Right: average frequencies of Foxp3^+^CD25^+^ Tregs. Right average numbers CD25^+^Foxp3^+^ in in vitro activated Tregs. (**O**) Line graph of calcium influx per time (s) analyzed by FlowJo Kinetics. Data from 2-3 independent experiments. Data shown as means ± SD. *p < 0.05, **p < 0.01, ***p < 0.001. Statistical significance was determined by unpaired t tests.

Subsequent analysis of T cell infiltration in the lungs and medLNs revealed increased numbers of CD44^+^CD8^+^ T cells in both organs (**Fig. S5A-B**). Though no numerical differences in Tregs were noted between WT and Treg-specific P2RX7-KO mice, higher numbers of CD4^+^ Teffs were present in the WT lungs (**Fig. 5E**). In the medLNs, increased numbers of both Tregs and CD4^+^ Teffs were observed in Treg-specific P2RX7-KO mice compared to WT counterparts. (**Fig. 5F**). Notably, an increased number of Live/dead^+^Foxp3^+^ Tregs were detected in both lungs and medLNs, suggesting higher cell death rates among P2RX7-KO Tregs (**Fig. 5G and S5C-D**).

To assess the suppressive capacity of tumor-infiltrating WT versus P2RX7-KO Tregs *ex vivo*, we sorted these cells from lung tumors and conducted suppression assays (**Fig. 5H and Fig. S1C**). Our data showed that WT lung Tregs were more potent in inhibiting the proliferation of *in vitro* activated CD4^+^ and CD8^+^ T cells compared to P2RX7-KO Tregs at 1:5 and 1:20 Treg:Teff ratios (**Fig. 5I-J**). This enhanced suppressive function correlated with higher expression of Ki-67, CD25, and CTLA-4 in WT Tregs (**Fig. 5K and S5E**). Conversely, in the medLNs, no differences were observed in Ki-67 and CTLA-4 expression, while CD25 was upregulated in P2RX7-KO Tregs (**Fig. 5L and S5F**). This data indicates a tissue-specific role of P2RX7 in promoting Treg function within the TME.

We next conducted *in vitro* experiments where Tregs were activated with anti-CD3/CD28 and either treated or not treated with the P2RX7-specific agonist HEI (HEI3090) (**Fig. 5M**) (28). Activation in the presence of HEI increased Treg numbers in culture (**Fig. 5N**). Given that P2RX7 functions as an ion channel (34), we performed a calcium influx assay, revealing that HEI-treated Tregs exhibited a higher basal calcium influx activity compared to control Tregs, which further increased upon ionomycin (Imy) addition. Additional HEI did not affect previously treated cells but enhanced calcium uptake in control Tregs. The P2RX7 antagonist A4 (A438079) reduced calcium influx only in control Tregs (**Fig. 5O**) (29). Collectively, these findings suggest that, despite the absence of numerical differences between WT and P2RX7-KO Tregs in this model, tamoxifen-induced ablation of P2RX7 diminishes the suppressive capacity of Tregs within lung tumors, possibly because of the reduction the P2RX7 action on calcium influx.

### Treg-intrinsic P2RX7 expression in Tregs limits type 1 and Tfh-like CD4^+^ T cell responses in lung cancer

In line with the lung tumor suppressive function of Tregs via P2RX7, we observed that tamoxifen-induced ablation of P2RX7 in Tregs enhances type 1 and cytotoxic responses in the lungs of tumor-bearing mice. Specifically, there was an increased accumulation of IFN-γ-producing CD4^+^ and CD8^+^ T cells (**Fig. 6A-B**), as well granzyme B-producing CD4^+^ and CD8^+^ T cells (**Fig. 6C-D**) in the lungs of Treg-specific P2RX7-KO mice compared to WT. Further phenotypic analysis of Teffs in WT and Treg-specific P2RX7-KO mice revealed a significant upregulation of markers characteristic of follicular helper T (Tfh) cells in the P2RX7-KO group. Heatmaps and gMFI analyses demonstrated higher expression of Bcl6, PD-1, CXCR5 and ICOS in Teff cells from both lungs and medLNs of Treg-specific P2RX7-KO mice compared to WT mice (**Fig. 6E-F**). Additionally, increased expression of the lung parenchyma homing marker CD69 (35) was noted in Treg-specific P2RX7-KO mice, while the lung vasculature-associated marker KLRG1 (35) was upregulated in CD4^+^ Teffs from WT mice (**Fig. 6E-F, S6A-D**), suggesting enhanced tissue localization of Teff cells within the lung parenchyma in the absence of P2RX7 in Tregs.

**Fig. 6.**
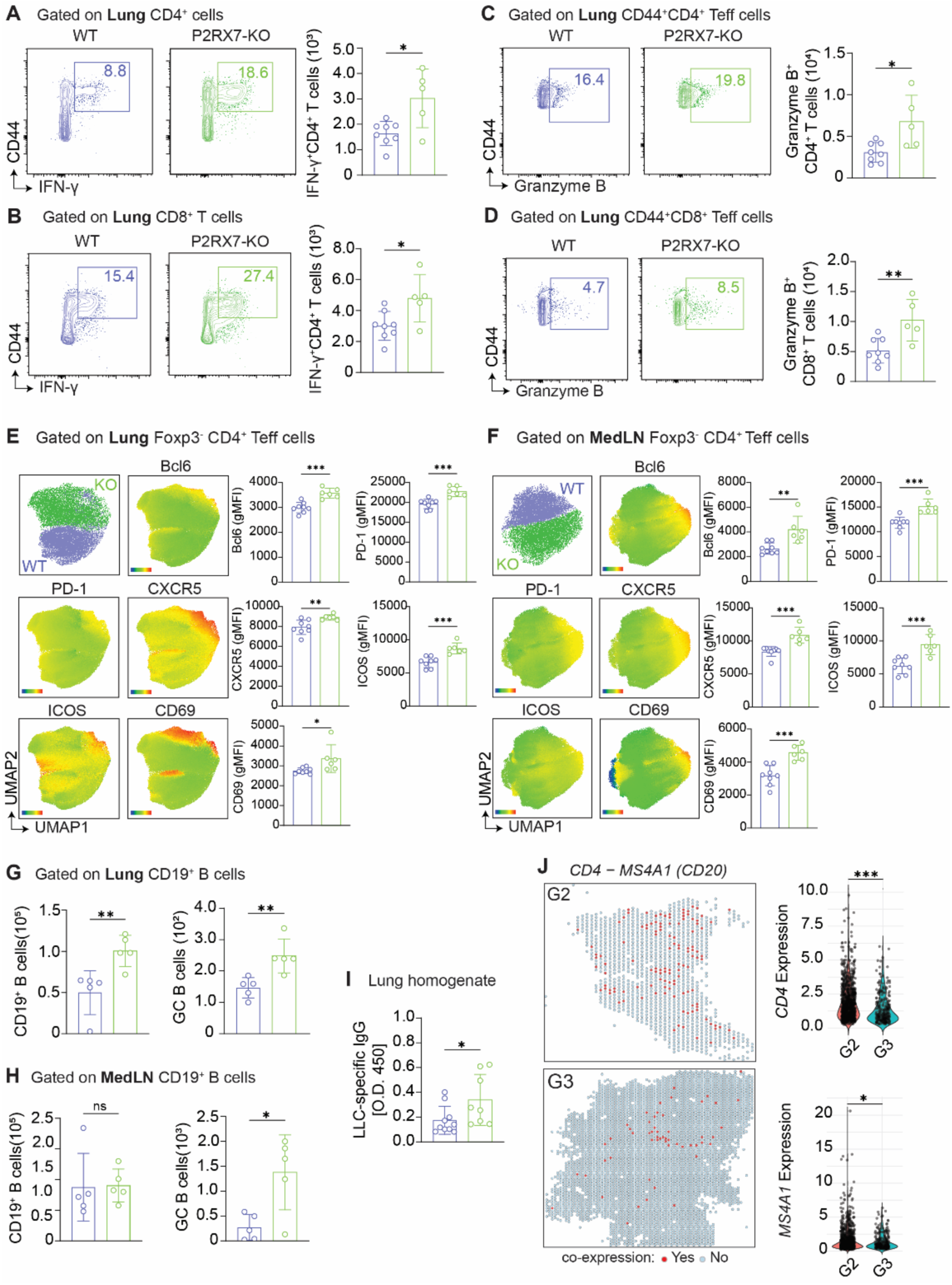
Effects of Treg-intrinsic P2RX7 on CD4^+^ T cell-mediated type 1 and Tfh-like responses in lung cancer experimental model. (**A-I**) WT (Foxp3 Cre-ERT2 *P2rx7^fl/fl^* - Vehicle) and Treg-specific P2RX7-KO (Foxp3 Cre-ERT2 *P2rx7 ^fl/fl^* – tamoxifen) mice were injected i.v. with LLC-GFP cells, and lung were collected on day 30 p.i. (**A**) Left: flow-cytometry plots of CD44 and IFN-γ expression on CD44^+^CD4^+^ RT cells in the lungs. Right: average numbers of IFN-γ-producing CD44^+^CD4^+^ T cells in the lungs. (**B**) Left: flow-cytometry plots of CD44 and IFN-γ expression on CD44^+^CD8^+^ T cells in the lungs. Right: average numbers of IFN-γ-producing CD44^+^CD8^+^ T cells in the lungs. (**C**) Left: flow-cytometry plots of CD44 and granzyme B expression on CD44^+^CD4^+^ RT cells in the lungs. Right: average numbers of granzyme B-producing CD44^+^CD4^+^ T cells in the lungs. (**D**) Left: flow-cytometry plots of CD44 and granzyme B expression on CD44^+^CD8^+^ T cells in the lungs. Right: average numbers of granzyme B-producing CD44^+^CD8^+^ T cells in the lungs. (**E**) Left: UMAP showing lung CD4^+^ Teff clusters, and heatmaps of Bcl6, PD-1, CXCR5, ICOS and CD69 expression in WT and Treg-specific P2RX7-KO mice. Right: average geometric mean fluorescence intensity (gMFI) of Bcl6, PD-1, CXCR5, ICOS and CD69 on lung Foxp3^-^CD44^+^CD4^+^ Teff cells. (**F**) Left: UMAP showing medLN CD4^+^ Teff clusters, and heatmaps of Bcl6, PD-1, CXCR5, ICOS and CD69 expression in WT and Treg-specific P2RX7-KO mice. Right: average geometric mean fluorescence intensity (gMFI) of Bcl6, PD-1, CXCR5, ICOS and CD69 on medLN Foxp3^-^CD44^+^CD4^+^ Teff cells. (**G**) Left: flow-cytometry plots of GL7 and Fas expression on CD19^+^ B cells in the lungs. Right: average numbers of GL7^+^Fas^+^ GC B cells in the lungs. (**H**Average numbers of GL7^+^Fas^+^ GC B cells in the medLNs. (**I**) Average optical density (O.D. 450 nm) of LLC-specific IgG in the lung homogenate. (**J**) Left: Spatial images showing the location of spots co-expressing *CD4* and *MS4A1* (CD20) in G2 and G3 lung cancer patients. Right: Violin plots showing the expression of *CD4* and *MS4A1* (CD20) in the spots co-expressing both genes in the tissue of G2 and G3 lung cancer patients. Data from 2-3 independent experiments. Data shown as means ± SD. *p < 0.05, **p < 0.01, ***p < 0.001. Statistical significance was determined by unpaired t tests.

Given the observed increase in Tfh-like cells in the lungs and medLNs of Treg-specific P2RX7-KO mice, we assessed the presence of germinal center (GC) B cells (GL7^+^Fas^+^CD19^+^) in these tumor-bearing mice. A notable increase in GC B cells was detected in both lungs and medLNs of P2RX7-KO mice compared to WT mice (**Fig. 6G-H**). Considering that interactions between Tfh and GC B cells in the lungs and lymph nodes lead to the production of antitumor antibodies (31, 21), we quantified LLC-specific IgG levels in the lung homogenates of tumor-bearing mice, and elevated levels of LLC-specific IgG were observed in P2RX7-KO mice. (**Fig. 6I**).

The interaction between GC B cells and Tfh-like cells in the lungs contributes to the formation of tertiary lymphoid structures (TLS) within tumors (8), facilitating antibody-mediated antitumor protection. We analyzed spatial transcriptomics data from patients with grade 2 and grade 3 lung cancer, focusing on spots co-expressing *CD4* and *MS4A1* (CD20, a B cell marker). While numerous spots were present in both groups, a higher number was observed in G2 patients (**Fig. 6K**). These findings suggest that interactions between CD4^+^ Teffs and B cells within lung tumors may be associated with better disease prognosis, mirroring observations from our lung cancer experimental model. Overall, our data supports the conclusion that P2RX7 enhances the suppressive activity of Tregs on type 1 and Tfh-like responses, thereby promoting tumor growth in the lung.

## DISCUSSION

eATP serves as a danger signal within the TME, signaling through purinergic receptors such as P2RX7 (17, 18, 36). While the eATP–P2RX7 axis has been studied across various cancers, including lung cancer, its specific role in T cells during lung tumor progression remained unclear (37). Our work demonstrated that P2RX7 expression in CD4⁺ T cells modulates the balance between Tregs and Teffs, thereby influencing tumor growth in the lung. Single-cell RNA sequencing analyses of advanced-stage

NSCLC patients revealed that *P2RX7* is highly expressed in T cells within the TME. Spatial transcriptomics further indicated that *P2RX7* expression is increased in G3 tumors compared to G2, suggesting its potential as a biomarker for advanced lung cancer stages and as a therapeutic target (38). Previous studies have shown that pharmacological inhibition or genetic ablation of P2RX7 can prevent tumor growth, potentially by promoting M1 macrophage polarization and enhancing tumor-infiltrating T cell responses (39). However, the specific impact of P2RX7 on T cells in lung cancer has not been fully elucidated. In melanoma models, P2RX7 limits the accumulation of CD4⁺ and CD8⁺ T cells within tumors by inducing their senescence, without affecting Treg accumulation (40). In our study, Tregs from advanced-stage NSCLC patients exhibited higher *P2RX7* expression compared to other CD4⁺ T cells. Spatial transcriptomics confirm increased co-expression of *P2RX7* and *FOXP3* in G3 tumors, increasing in higher disease stages. Similarly, in a murine lung cancer metastasis model, tumor-infiltrating Tregs express higher levels of P2RX7 than Teffs. Using various experimental models, we demonstrated that P2RX7 expression in CD4⁺ T cells promotes lung tumor growth, whereas its ablation enhances Teff accumulation and reduces tumor burden. Previous studies from our group have demonstrated that the eATP–P2RX7 axis is crucial for the accumulation and localization of CD4⁺ T cells in response to severe tuberculosis and influenza infections (22, 41). Therefore, the impact of P2RX7 on T cell responses to tumors may vary depending on the tumor site and/or the type of malignancy.

The increased Teff numbers in T cell-specific P2RX7-KO mice may result from reduced Treg accumulation in the lungs. *In vitro* activation of Tregs in the presence of a P2RX7 agonist (HEI3090) mimics the Treg expansion observed in WT mice. Contrastingly, previous studies using BzATP, a direct P2RX7 agonist, reported Treg apoptosis (42). The use of HEI3090, which sensitizes P2RX7 to ATP without directly activating it, may explain these differing outcomes by enhancing receptor sensitivity to autocrine ATP release (28).

Treg accumulation in lung cancer impairs effective CD4⁺ and CD8⁺ Teff responses and compromises immunotherapy efficacy (11). In Treg-specific P2RX7-KO mice, we observed that while Treg numbers remain unchanged, however their suppressive functions are diminished. Tamoxifen-induced P2RX7 ablation in Tregs leads to decreased expression of Ki-67, CTLA-4, and CD25, markers associated with Treg proliferation and suppressive capacity (43, 44). Given that T cell receptor (TCR) activation and calcium influx are crucial for upregulating these molecules (45) and considering the role of P2RX7 as an ion channel facilitating calcium entry (34), our findings suggest that P2RX7 signaling may enhance Treg suppressive function via calcium-mediated pathways. Future research will be necessary to formally establish this connection between P2RX7, calcium influx and Treg suppression. We also found that, while Treg numbers increase in the medLNs of P2RX7-KO mice, the suppressive markers reduced in lung Tregs are not similarly affected in medLN Tregs. This indicated that the influence of P2RX7 on Treg function is tissue-specific, primarily affecting Tregs upon their arrival in the lung tissue. The molecular cues driving this tissue-specific regulation warrant further investigation.

In the absence of Treg-mediated suppression through cell-specific P2RX7 ablation, the protective functions of CD4⁺ and CD8⁺ Teffs are increased. Reduced Treg suppression in P2RX7-KO mice enhances type 1 Teff responses, characterized by increased IFN-γ and granzyme B production, key antitumor mediators (46, 47). The most pronounced effects are observed in Tfh-like cells, which are primary Treg targets due to their high PD-1 expression (48, 49). P2RX7-KO Tregs fail to regulate Teffs effectively, leading to Tfh-like cell accumulation in lung tumors and medLNs. This accumulation promotes TLS formation within tumors and germinal centers development in lymph nodes, enhancing antibody-mediated tumor cell elimination (8). Consequently, we observed increased production of tumor-specific IgG and reduced tumor growth in the presence of P2RX7-KO Tregs. The presence of TLS in lung cancer patients correlates with favorable prognosis (8), and our spatial transcriptomics data show higher CD4⁺ and B cell co-localization in G2 patients. In summary, our findings suggested that in response to lung cancer, P2RX7 promoted Treg infiltration and immunosuppression, limiting Teff accumulation and both type 1 and humoral immune responses. Targeting P2RX7 may offer a therapeutic strategy to mitigate Treg-mediated suppression and enhance Teff-mediated protection against lung cancer.

## ACKNOWLEDGMENTS

We thank the Borges da Silva, Lancaster and D’Imperio-Lima labs for their intellectual support. We also thank the Mayo Clinic Arizona Flow Cytometry and Histology Core for experimental support.

