## Supplementary Data for "Sensing of extracellular ATP via P2RX7 drives lung tumor growth through regulatory T cell suppressive function"

#### SUPPLEMENTARY MATERIAL

### Sensing of extracellular ATP via P2RX7 drives lung tumor growth promoting regulatory T cell accumulation and suppression

**Authors:** Igor Santiago-Carvalho[1†], Ronaldo Francisco Jr[2], Bruna de Gois Macedo[1], Caio Loureiro Salgado[1], Carly R. Stoll[1], Marcos Pinheiro Cione[1,3], Emily White[1], Tyler Johnston[1], Chloe Liliana Leff[1,5], Ildefonso Silva Junior[1], Fabio Carvalho de Souza[1], Maria Regina D'Império Lima[3], Jessica Naomi Lancaster[1] and Henrique Borges da Silva[1,4†]

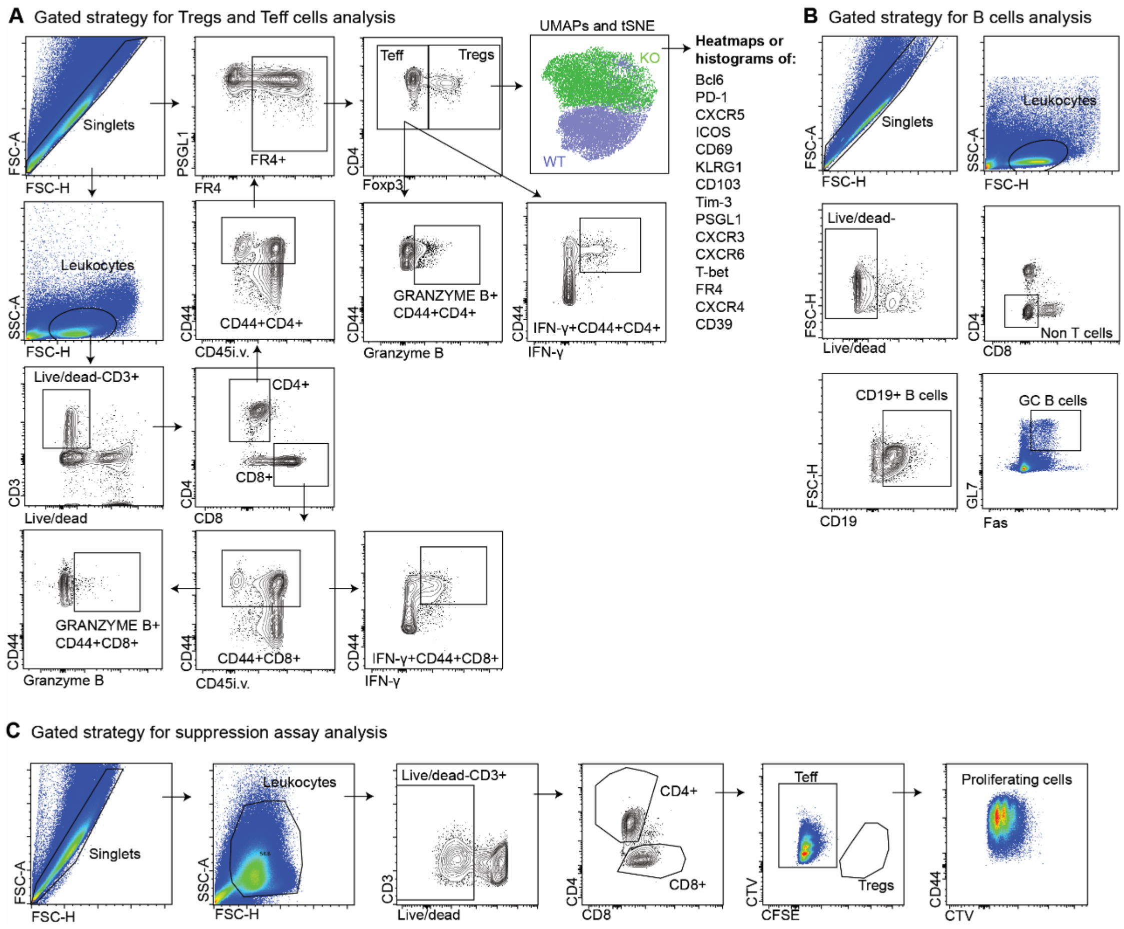


**Fig. S1. Gate strategy for flow cytometry analysis.** (**A**) Gate strategy for analysis of T cells in the lungs, medLNs and spleens of mice with lung cancer. (**B**) Gate strategy for analysis of B cells in the lungs and medLNs of mice with lung cancer. (**C**) Gate strategy for analysis of *ex vivo* Tregs suppression assay.

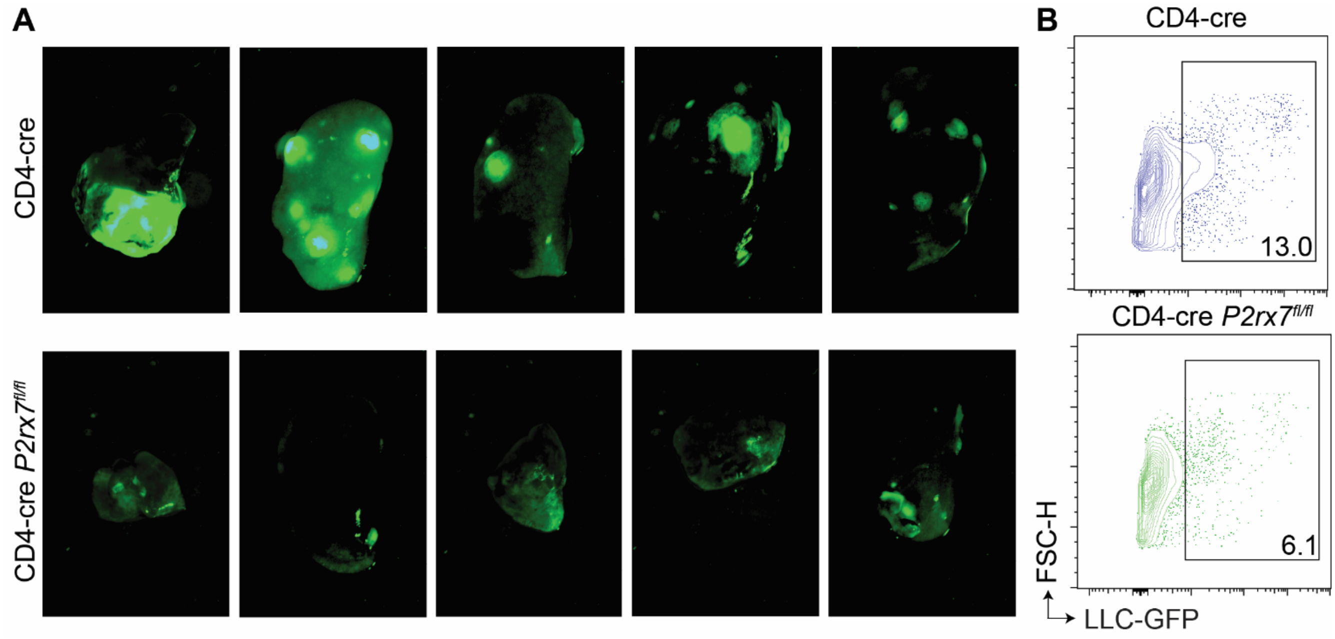


**Fig. S2. Effects of T-cell intrinsic P2RX7 on the growth of tumors in the lung.** WT (CD4-cre) and T cell-P2RX7-KO (CD4-cre *P2rx7*$^{fl/fl}$) mice were injected i.v. with LLC-GFP cells and lungs were collected on day 30 p.i. (**A**) Representative fluorescence macroscopy of the left upper lung lobes with tumors (20 μm). (B) Flow-cytometry plots of GFP expression on LLC cells in the lungs of WT and T cell-P2RX7-KO mice.

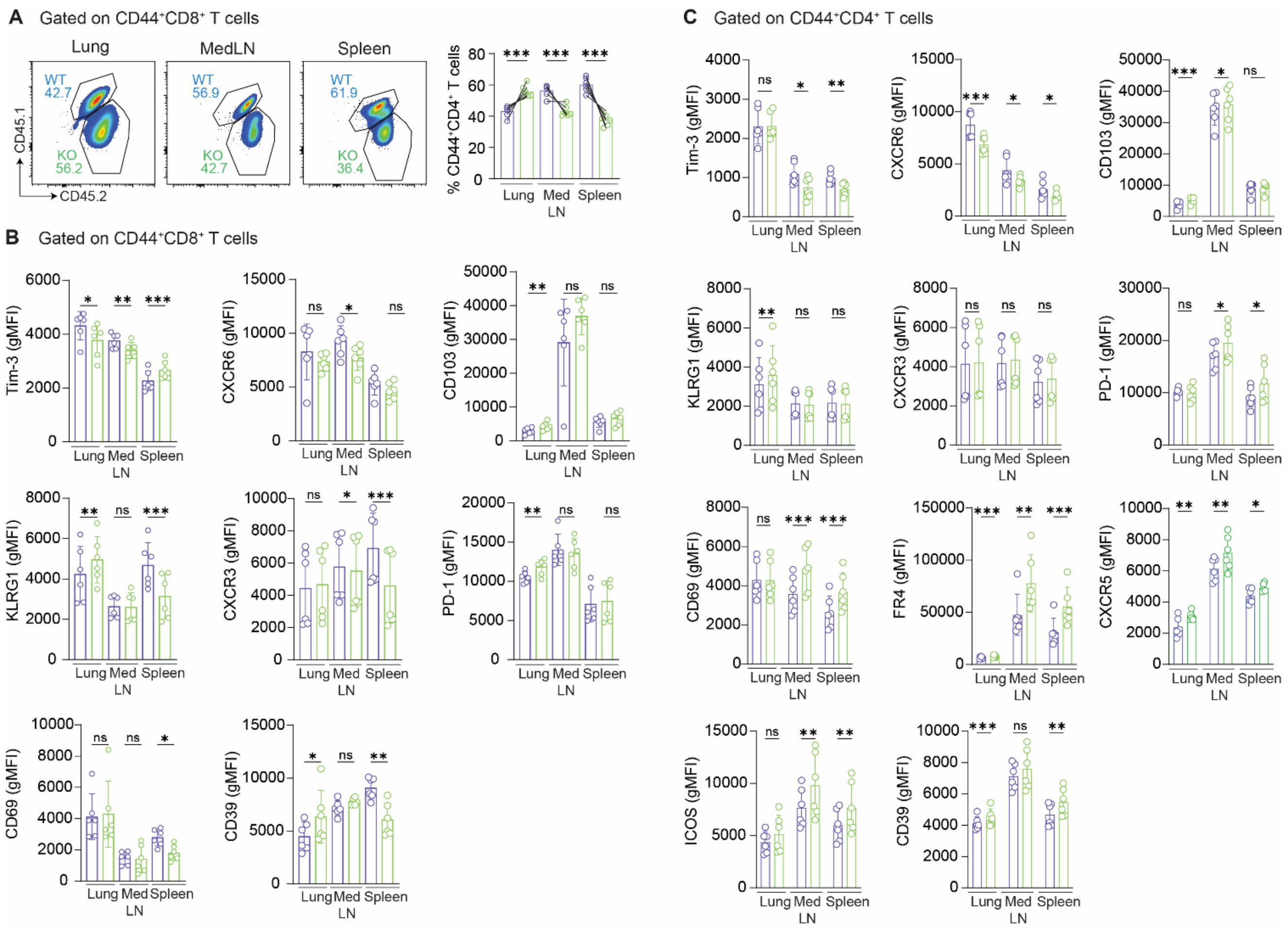


**Fig. S3. Effects of P2RX7 expression on T cells phenotype in lung cancer experimental model.** RAG-KO mice were reconstituted with splenocytes from WT and P2RX7-KO mice and injected i.v. with LLC-GFP cells. On day 30 p.i. the lungs, medLNs and spleens were collected. (**A**) Left: flow-cytometry plots of CD45.1 and CD45.2 on CD44$^+$CD8$^+$ T cells. Right: average frequencies and numbers of CD45.1 and CD42.2 CD44$^+$CD8$^+$ T cells. (**B**) Average geometric mean fluorescence intensity (gMFI) of Tim-3, KLRG1, CXCR3, CXCR6, CD39, CD69, PD-1 and CD103 on CD44$^+$CD8$^+$ T cells. (**C**) Average geometric mean fluorescence intensity (gMFI) of PD-1, CD103, Tim-3, KLRG1, CXCR3, CXCR6, ICOS, CD69, FR4 and CD39 on CD44$^+$CD8$^+$ T cells. Data are from 2-3 independent experiments. Data shown as means ± SD. $^*p < 0.05$, $^{**}p < 0.01$, $^{***}p < 0.001$. Statistical significance was determined paired t tests.

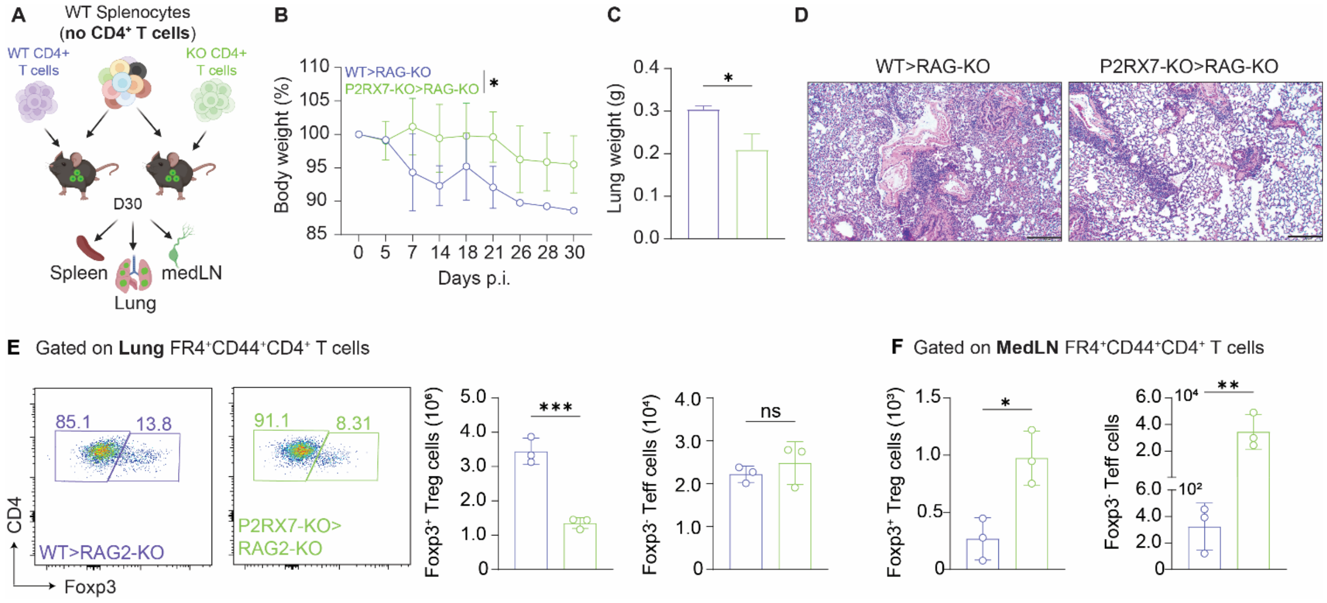


**Fig. S4. Effects of CD4$^+$ T-cell intrinsic P2RX7 on tumor growth and Treg accumulation in the lungs.** RAG2-KO mice were transferred with CD4$^+$ depleted splenocytes from WT mice and then were randomly assigned into two groups, receiving either CD4$^+$ T cells from WT (WT>RAG2-KO) or P2RX7-KO (P2RX7-KO>RAG2-KO). 24 hours later, both groups were injected intravenously with LLC-GFP tumor cells. (**A**) Schematic illustration showing the lung cancer experimental protocol (BioRender.com). (**B**) Average body weights (percentages related to day 0). (**C**) Lung weight values on day 30 p.i. (**D**) Representative section of the left upper lung lobes stained with H&E (200 µm). (**E**) Left: flow-cytometry plots of CD4 and Foxp3 expression in the lungs. Right: average numbers of CD44$^+$CD4$^+$ Tregs (Foxp3$^+$) and Teffs (Foxp3$^-$) in the lungs. (**F**) Average numbers of CD44$^+$CD4$^+$ Tregs (Foxp3$^+$) and Teffs (Foxp3$^-$) in the medLNs. Data are from 2-3 independent experiments. Data shown as means ± SD. *p < 0.05, **p < 0.01, ***p < 0.001. Statistical significance was determined by unpaired t tests.

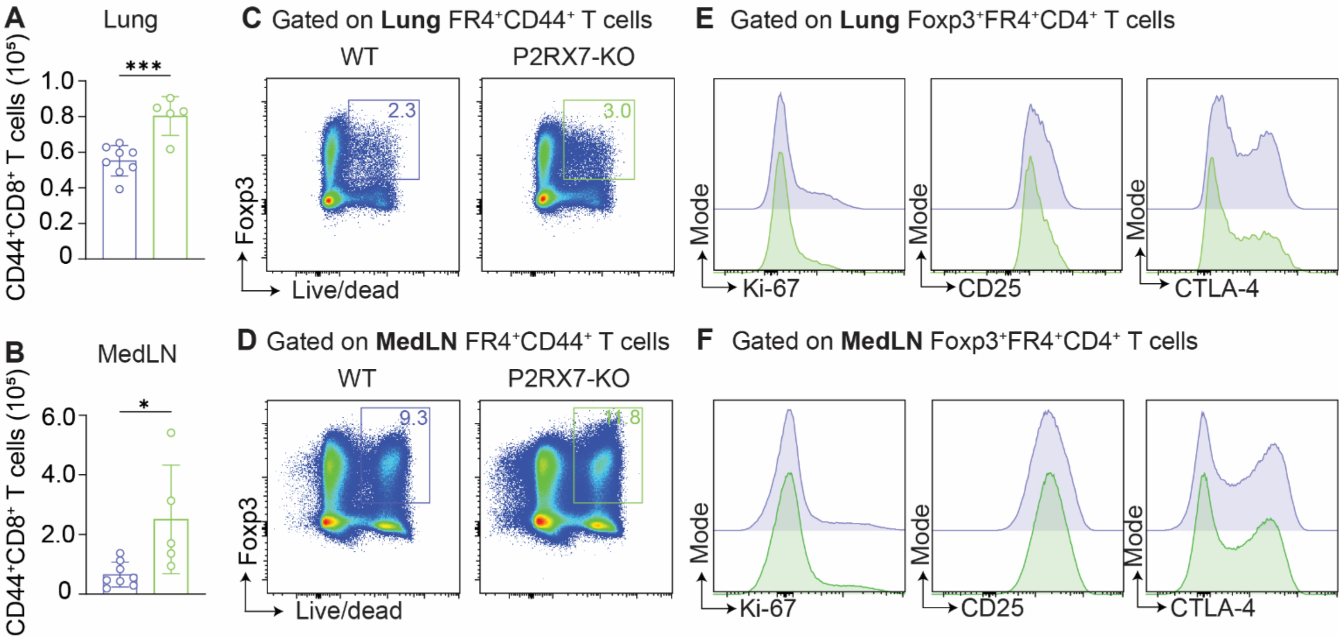


**Fig. S5. Effects of P2RX7 expression on Tregs in lung cancer experimental model.** WT (Foxp3 Cre-ERT2 *P2rx7*$^{fl/fl}$ - Vehicle) and Treg-specific P2RX7-KO (Foxp3 Cre-ERT2 *P2rx7* $^{fl/fl}$ – TAM) mice were injected i.v. with LLC-GFP cells and lung were collected on day 30 p.i. (**A**) Average numbers of $CD44^+CD8^+$ T cells in the lungs. (**B**) Average numbers of $CD44^+CD8^+$ T cells in the medLNs. (**C**) Flow-cytometry plots of Foxp3 and Live/dead expression in the lungs. (**D**) Flow-cytometry plots of Foxp3 and Live/dead expression in the medLNs. (**E**) Histograms of Ki-67, CD25, and CTLA-4 expression in the lungs. (**F**) Histograms of Ki-67, CD25, and CTLA-4 expression in the medLNs. Data are from 2-3 independent experiments. Data shown as means ± SD. *p < 0.05, **p < 0.01, ***p < 0.001. Statistical significance was determined by unpaired t tests.

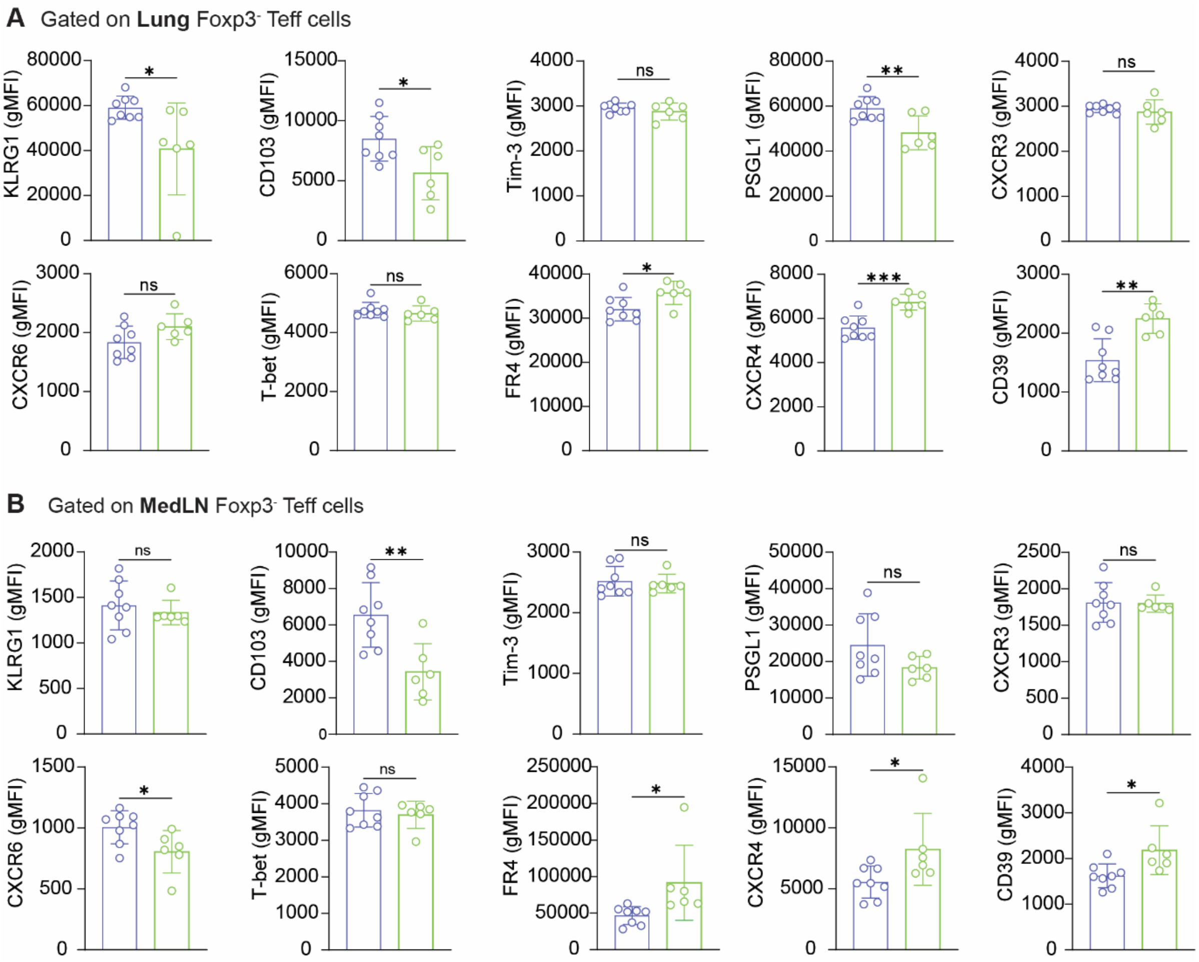


**Fig. S6. Effects of Treg-specific P2RX7 ablation on Teffs in experimental lung cancer model.** WT (Foxp3 Cre-ERT2 *P2rx7$^{fl/fl}$* - Vehicle) and Treg-specific P2RX7-KO (Foxp3 Cre-ERT2 *P2rx7 $^{fl/fl}$* – TAM) mice were injected i.v. with LLC-GFP cells, and lung were collected on day 30 p.i. (**A**) Average geometric mean fluorescence intensity (gMFI) of KLRG1, CD103, Tim-3, PSGL1, CXCR3, CXCR6, T-bet, FR4, CXCR4 and CD39 on lung $Foxp3^-CD44^+CD4^+$ Teffs. (**B**) Average geometric mean fluorescence intensity (gMFI) of KLRG1, CD103, Tim-3, PSGL1, CXCR3, CXCR6, T-bet, FR4, CXCR4 and CD39 on medLN $Foxp3^-CD44^+CD4^+$ Teffs. Data are from 2-3 independent experiments. Data shown as means ± SD. *$p < 0.05$, **$p < 0.01$, ***$p < 0.001$. Statistical significance was determined by unpaired t tests.
